# Uninterpretable graphs at a scientific conference

**DOI:** 10.1101/2024.11.08.622642

**Authors:** Zen Faulkes

## Abstract

Comparing averages requires knowing sample variation, which can be shown visually using a bar graph with error bars or a box plot. Because error bars and box plots can show different measures, they must be labelled to be interpretable. Of graphs of averages shown at a scientific conference in 2020, 87.5% of graphs did not contain enough information on the slide to interpret them. The only graphs that were fully interpretable (12.5%) were bar graphs. By comparison, 92% of graphs of averages published in a journal in the same field contained enough information to interpret them. While conferences are not and should not be as stringent about data presentation as journals, that a statistical value as simple as an average are not clearly presented suggests graphs are created by rote.

## Introduction

Averages are simple and widely recognized statistical measurements that are fundamental to many scientific analyses. But interpreting an average requires knowing the distribution and range of numbers that was used to calculate a mean or median. In scientific graphs, this variation is shown with error bars in bar graphs or by drawing range bars (Spear 1952) or box plots (McGill et al. 1978, Tukey 1970). Summary statistics shown in bar graphs are often criticized because they can conceal interesting patterns in the data (Anscombe 1973, Matejka and Fitzmaurice 2017, Weissgerber et al. 2015). Data displays like box plots (McGill et al. 1978, Spear 1952, Tukey 1970) can show central tendency and data distribution in more detail.

But just as “average” can be measured multiple ways (e.g., mean, median, mode), error bars and box plots can show variation in multiple ways. Error bars in bar graphs can show standard deviation, standard error of the mean, confidence interval, or range (minimum and maximum) (Cumming et al. 2007). For a given data set, standard deviation, standard error, and confidence interval can vary widely (Figure 1). Further, these measures are affected by sample size in different ways: standard error and confidence intervals become smaller as sample size increases, while standard deviation does not (Cumming et al. 2007).

**Figure 1.**
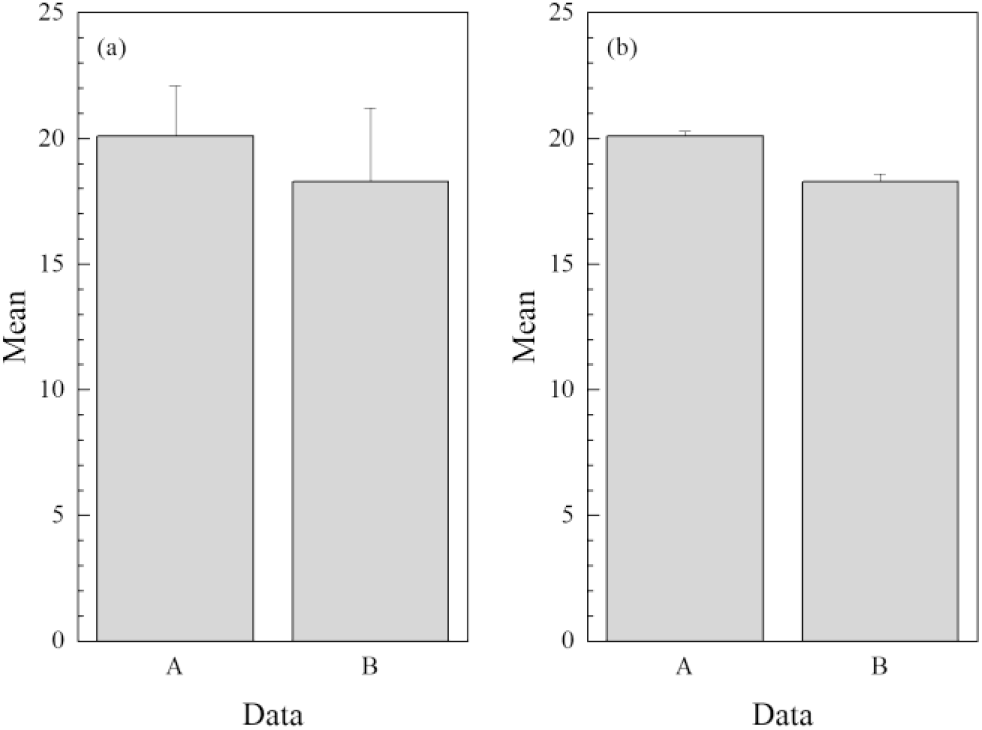
(a) Means of two randomly generated data sets (n=100). Error bars show standard deviations. sample a was drawn from a normally distributed population with a mean is 20 and standard deviation of 2; sample b was drawn from a normally distributed population with means of 18 and a standard deviation of 3. (b) Same data as (a), except error bars show standard error.

Similarly, box plots can show data in many ways. A technical graphic software package, Origin 2020 (OriginLab corporation), offers many options for creating elements in box plots: two options for the central line (median and mean), six options for the box (including “custom”) and ten options for the whiskers (including “custom”).

Regardless of whether a bar graph or box plot is used to display averages, it is critical that there is enough information about what is plotted to interpret the data. Graphs increase persuasiveness of a claim (Tal and Wansink 2016), and the length of error bars can change the impression of an average, particularly when averages are compared. Regardless of statistical tests, many researchers shown the graphs in Figure 1 might differ in how they judged whether the difference between the two groups was “real” depending on which graph they saw, even though the data are the same. Researchers may choose measures of variation based on “whatever is smallest” to instill more confidence in the viewer.

A bar graph should show a mean value, error bars, and identify the measure shown by the error bars. A box plot should show a median value, central quartiles (i.e., 50% of the data), and identify the measure shown by the whiskers. Additional detail can be added by showing a calculated data distribution (e.g., violin plot) or the entire raw data set in addition the summary statistics (Weissgerber et al. 2015).

These issues of data display should be widely known in research communities, but are they followed in practice? Scientific conferences provide a useful window into the work habits of scientists regarding data display. Data presented at scientific conferences rarely undergo the same level of scrutiny as journal articles, where peer review and editorial processes should ensure that the elements required to interpret graphs are included in published articles. Because scientists follow common practices in their field, it is useful to investigate whether data displays in conferences show best practices of data display. I analyzed graphs from the 2020 meeting of the Animal Behavior Society and the journal *Animal Behaviour* to test how often best practices for data display are followed.

## Methods

The virtual Animal Behavior Society meeting was held online from 27-31 July 2020 and contained 900 recorded presentations. I haphazardly sampled presentations recorded for the Animal Behavior Society and looked for data slides that included a graph that showed some form of average. I omitted any presentations that included a request not to share the presentation on social media. If I found a graph displaying an average, I captured an image of the slide and recorded the talk title and URL. All recorded sessions from all days of the conference were sampled. I continued this process until I reached 200 images from 200 separate presentations.

For comparison, I examined all figures printed in volume 165 of the journal *Animal Behaviour*, which had a cover date of July 2020 (that is, the same month as the conference). I scored each graph in a compound figure (e.g., Figure 1a, 1b, 1c) independently. Volume 165 contained 52 figures that displayed averages.

I categorized each graph sampled from the conference or published in the journal as a bar graph or box plot. I categorized a graph as a bar graph if it showed a single value of central tendency, regardless of whether that value was shown as a bar or a dot. I examined bar graphs for error bars and any indication on the slide of what the error bar showed (standard deviation, standard error, confidence interval). I also recorded whether the raw data or distribution of data were shown in addition to the mean. I categorized a graph as a box plot if it showed a rectangle divided by a central line with whiskers emerging from opposite sides. I examined box plots for any indication of the components of the graph where, and whether the raw data or distribution of data were shown in addition to the box.

## Results

Conference graphs were almost evenly split between bar graphs (53%, 106/200) and box plots (47%, 94/200).

Most conference bar graphs had error bars (98%, 104/106), but most of those (76%, 79/104) did not show on the slide what the error bars represented. Of bar graphs that did indicate what the error bars were (24%, 25/104), three different measures of variance were represented: standard error (60%, 15/25), standard deviation (16%, 4/25), and confidence interval (24%, 6/25). Few bar graphs were presented with raw data (9%, 10/106) or data distributions (1%, 1/106).

No box plots (0%, 0/94) were labelled in a way that indicated all three components. Only one graph indicated the horizontal line represented the median and that the box showed the interquartile range (“IQR” on the graph), but what the whiskers represented were not indicated. More box plots (41%, 39/94) showed the raw data than bar graphs (9%). Like bar graphs, few box plots (2%, 2/94) showed data distributions.

The journal *Animal Behaviour* had more bar graphs (73%, 38/52) than box plots (27%, 14/52). Almost all journal graphs (92%, 48/52) contained complete information, whether presented as bar graphs (29/52) or box plots (14/52). The graphs that did not completely specify their elements were preceded in their paper by similar graphs: i.e., Figure 2 described its error bars, but Figure 5 did not. Because the two figures share the same format, it is reasonable to assume that both use error bars the same way.

**Figure 2.**
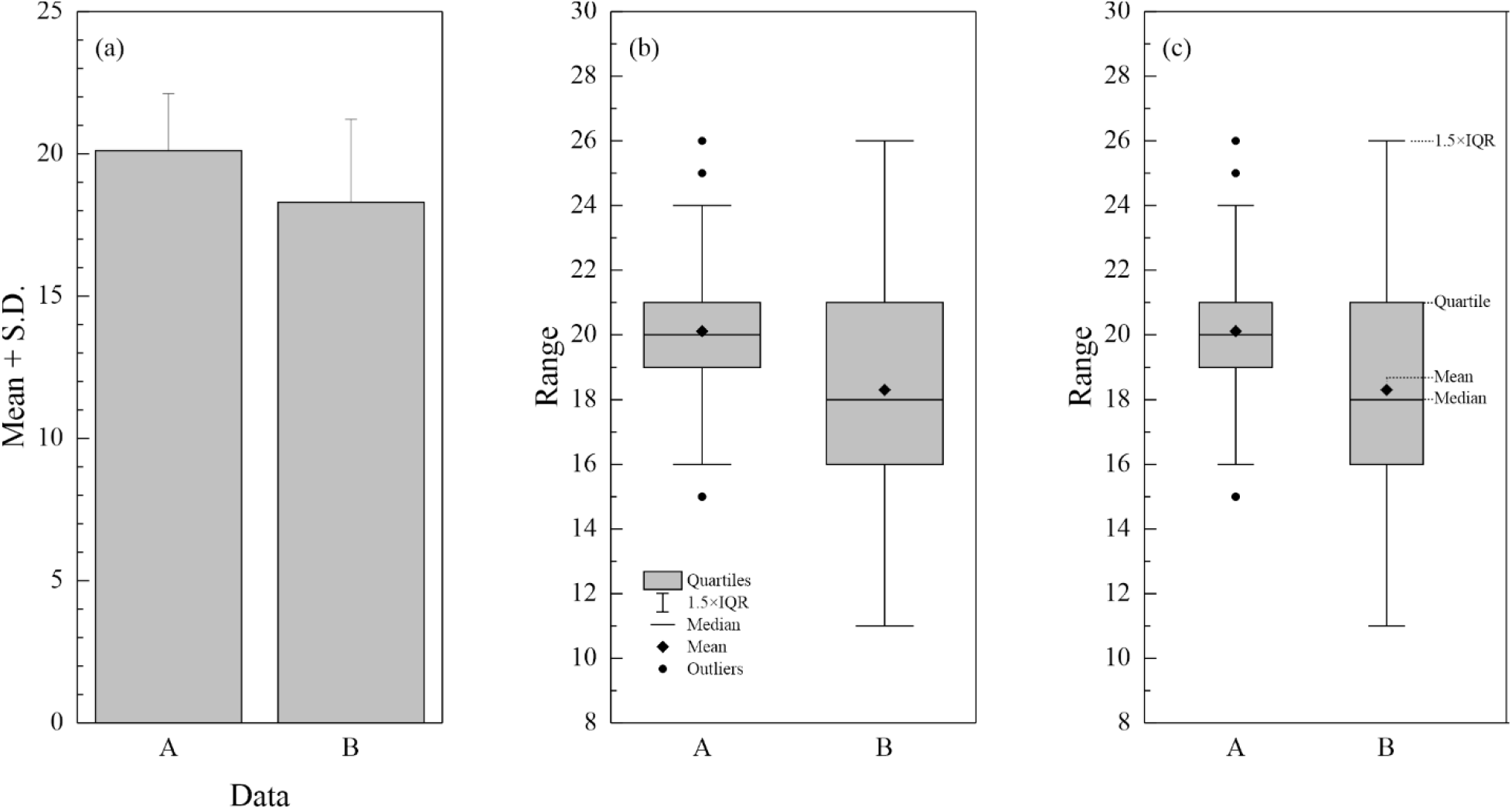
(a) Bar graph with labelled Y axis. (b) Box plot showing component descriptors in legend. (c) Box plot with labelled component descriptions.

Error bars most often showed standard error (97%, 33/34), but standard deviation was also displayed (3%, 1/34). The horizonal line of box plots all showed the median, and the boxes all showed quartiles above and below the media (i.e., 50% of data). Whiskers, however, were not consistent in their use. Some (57%, 8/14) showed the range of data (i.e., minimum and maximum values) while others (43%, 6/14) showed a multiple of quartile values (1.5 × quartile). The information on error bars or box plot components was usually contained in the figure legend (90%, 43/48) rather than in the axes of the graph (10%, 5/48).

## Discussion

Data displays presented in a conference rarely followed best practices. Only 12.5% of graphs examined from an Animal Behavior Society conference included all the information necessary to interpret the averages shown, compared to at least 92% of published graphs. Conference graphs frequently left out what the error bars in bar graphs show and always left out what whiskers in box plots show. Box plots, often described as a more transparent way of showing data in theory, may be less transparent in practice because people take less care in describing or labelling them.

Published graphs show that the information encoded by these elements is not standardized. What error bars, boxes, and whiskers show on a graph cannot be assumed but must be provided explicitly by the presenter or author. Conference presenters may describe these elements verbally in their pre-recorded talks, but there are at least two problems with this. First, a good graphic design principle to put information at the point of need (Faulkes 2021) by integrating data and text (Tufte 2006). In a conference slide, labels should be on the slide itself so that they are visible when the data are visible. Second, describing these features verbally makes this critical information inaccessible to people who cannot hear the recorded commentary if closed captions are not provided.

In most cases, bar graphs can describe the error bars in the Y axis with annotations such as “Mean + SD” for standard deviation, “SE” or “SEM” for standard error, or “CI” for confidence interval (Figure 2a). Box plots have several elements that need description which will probably not fit in many axes. These might be listed in a legend (Figure 2b) or the elements might be labelled directly on the rightmost box (Figure 2c).

Conference presentations are a critical point to show best practices, because many researchers adopt the common practices of their field. It is possible, for example, that one reason that conference graphs do not describe graph features directly within the graph, is because journals often put the information in a figure legend. This may lead to presenters ignoring the information needed in their presentation slides.

That so many researchers do not provide enough information to interpret their graphs suggests researchers are creating graphs by rote, and that graphs and statistical analyses generally are used to create a superficial appearance of professional diligence in analysis.

## Supporting information

S1. Screenshots of conference data slides analyzed in this paper. https://figshare.com/s/dde317121d572a4d455e

S2. Spreadsheet of data analyzing graphics in conference and journal. https://figshare.com/s/570dcbd0bfd08b3359c3

